# Multiple Independent Genetic Code Reassignments of the UAG Stop Codon in Phyllopharyngean Ciliates

**DOI:** 10.1101/2024.07.18.603678

**Authors:** Jamie McGowan, Thomas A. Richards, Neil Hall, David Swarbreck

**Affiliations:** Earlham Institute, Norwich Research Park, Norwich, NR4 7UZ, UK; Department of Biology, University of Oxford, Oxford, OX1 3SZ, UK; School of Biological Sciences, University of East Anglia, Norwich, NR4 7TJ, UK

**Keywords:** Genetic code, stop codon reassignment, suppressor tRNA, Phyllopharyngea, Ciliophora, codon usage, phylogenomics

## Abstract

The translation of nucleotide sequences into amino acid sequences, governed by the genetic code, is one of the most conserved features of molecular biology. The standard genetic code, which uses 61 sense codons to encode one of the 20 standard amino acids and 3 stop codons (UAA, UAG, and UGA) to terminate translation, is used by most extant organisms. The protistan phyla Ciliophora (the ‘ciliates’) are an unusual exception to this norm, exhibiting the greatest diversity of non-canonical nuclear genetic code variants and evidence of repeated changes in code. In this study, we report the discovery of multiple independent genetic code changes within the Phyllopharyngea class of ciliates. By mining publicly available ciliate genome datasets, we discovered that three ciliate species from the TARA Oceans eukaryotic metagenome dataset use the UAG codon to putatively encode leucine. We identified novel suppressor tRNA genes in two of these genomes. Phylogenomics analysis revealed that these three uncultivated taxa form a monophyletic lineage within the Phyllopharyngea class. Expanding our analysis by reassembling published phyllopharyngean genome datasets led to the discovery that the UAG codon had also been reassigned to putatively code for glutamine in *Hartmannula sinica* and *Trochilia petrani*. Phylogenomics analysis suggests that this occurred via two independent genetic code change events. These data demonstrate that the reassigned UAG codons have widespread usage as sense codons within the phyllopharyngean ciliates. Furthermore, we show that the function of UAA is firmly fixed as the preferred stop codon. These findings shed light on the evolvability of the genetic code in understudied microbial eukaryotes.

## Introduction

Ciliates are a diverse phylum of single-celled eukaryotes (protists) characterised by their unusual genome biology. Interestingly, ciliate species exhibit the greatest diversity of non-canonical nuclear genetic codes, with several ciliate lineages possessing genetic codes which deviate from the standard genetic code. The standard genetic code uses three stop codons (UAA, UAG, and UGA) to terminate translation and 61 sense codons to encode an amino acid (Crick 1968). Once thought to be universal as it is used by most extant organisms, the standard genetic code is one of the most conserved features of molecular biology, emerging prior to the last universal common ancestor (LUCA) (Keeling 2016).

All reported genetic code changes in ciliates involve the reassignment of one or more stop codons to encode an amino acid. The most common non-canonical genetic code in ciliates (and eukaryotes in general) involves the reassignment of both the UAA and UAG (i.e., UAR) codons to encode glutamine, as observed in several lineages of ciliates, including the model ciliate species *Tetrahymena thermophila* and *Oxytricha trifallax* (Lozupone et al. 2001). In comparison, the UAR codons are reassigned to glutamic acid in the peritrichs (Wang et al. 2021) and to tyrosine in the *Mesodinium* genus (Heaphy et al. 2016). Whereas the UGA stop codon has been reassigned to cysteine in *Euplotes* (Meyer et al. 1991) and to tryptophan in *Blepharisma* (Lozupone et al. 2001). In some lineages of ciliates, including Karyorelictea and *Condylostoma*, all three stop codons have been reassigned and can have dual meanings encoding an amino acid or terminating translation depending on their context (Heaphy et al. 2016; Swart et al. 2016; Seah et al. 2022). In most reported non-canonical genetic code changes, the codons UAA and UAG have the same meaning, either they are both reassigned to code for an amino acid or they are both retained as stop codons, suggesting that evolutionary or mechanistic constraints couple the function of these two codons (Kollmar and Mühlhausen 2017; Pánek et al. 2017).

In this study, we report the discovery of three independent genetic code change events within the Phyllopharyngea class of ciliates. The Phyllopharyngea class is relatively understudied compared to other ciliate groups and includes taxa with diverse morphologies and lifestyles, including free-living species and symbiotic species (Lynn 2010). They include some of the most destructive parasites of fish (Bastos Gomes et al. 2017). Mining publicly available ciliate genome sequences, we identified three uncultivated ciliate species from the TARA Oceans eukaryotic metagenome dataset (Delmont et al. 2022) where the UAG stop codon has been reassigned to code for leucine. We identified novel suppressor tRNA genes with CUA anticodons in two of these genomes which are predicted to decode the reassigned UAG codon to leucine. Phylogenomic analysis revealed that these three uncultivated taxa form a monophyletic lineage within the Phyllopharyngea class. Reassembly and annotation of seven other phyllopharyngean genome sequences from previously published datasets (Maurer-Alcalá et al. 2018; Pan et al. 2019) revealed that *Hartmannula sinica* and *Trochilia petrani* have also undergone genetic code reassignments where UAG has been reassigned to encode glutamine. The other five phyllopharyngean species use the standard genetic code. Phylogenomic analysis infer that the reassignment of the UAG stop codon to encode glutamine has evolved independently twice within the Phyllopharyngea lineage. Thus, Phyllopharyngea contains a mix of species that use canonical and non-canonical genetic codes, with at least three independent genetic code change events. The reassigned UAG codon demonstrates widespread usage in the predicted proteomes as a sense codon in all five species showing reassignment, suggesting that its function is fixed as a sense codon. Furthermore, these data demonstrate that UAA is ubiquitously used as the preferred stop codon in these taxa and is therefore unlikely to later be reassigned as a sense codon. These findings reveal further divergences between the function of the UAA and UAG codons signifying repeat breaking of the proposed mechanistic constraints linking the function of UAA and UAG codons.

## Results

### Reassignment of the UAG stop codon to leucine in an uncultivated lineage of ciliates

To investigate the evolution of the genetic code in uncultivated ciliates, we mined eukaryotic metagenome-assembled genomes (MAGs) from the TARA Oceans project (Delmont et al. 2022). This is a dataset of manually curated genome assemblies. 30 MAGs from this dataset were classified as being ciliates, which we analysed in this study. Two complementary tools were initially used to predict the genetic code of each genome assembly – Codetta and PhyloFisher (Tice et al. 2021; Shulgina and Eddy 2023). Codetta predicts the meaning of each codon by aligning hidden Markov models from the Pfam database against a six-frame translation of a query genome assembly. The “genetic_code_examiner” utility from PhyloFisher predicts the genetic code by identifying and counting codons that correspond to highly conserved amino acid sites in a database of orthologous proteins.

This analysis revealed a novel genetic code change in ciliates. Three of the TARA Oceans ciliate MAGs were predicted to have reassigned the meaning of the UAG codon to encode leucine – TARA_ARC_108_MAG_00274, TARA_ARC_108_MAG_00306, and TARA_SOC_28_MAG_00066. We will refer to these as ARC_274, ARC_306, and SOC_66, respectively hereafter. Two of these MAGs are from the Arctic Ocean (ARC_274 and ARC_306), and the other MAG is from the Southern Ocean (SOC_66) (Delmont et al. 2022). ARC_306 (33.6 Mb; 77.8% BUSCO completeness) and SOC_66 (21.3 Mb; 49.8% BUSCO completeness) both have reasonable estimated genome completeness (**Table S1**). The third MAG ARC_274 has low completeness (11 Mb; 12.9% BUSCO completeness) which is reflected by its smaller assembly size (**Table S1**). Codetta identified 1011, 1898, and 2583 UAGs codons with a Pfam alignment in ARC_274, ARC_306, and SOC_66, respectively, and predicted that all three MAGs translate UAG to leucine with a log decoding probability of zero (**Fig. S1**). From the PhyloFisher dataset of 240 eukaryotic orthologs, PhyloFisher identified 8, 12, and 44 internal UAGs codons in ARC_274, ARC_306, and SOC_66, respectively, which correspond to positions where leucine is highly conserved (> 70% conservation) (**Fig. S2**). This is a very strict analysis as it only considers amino acid positions that are highly conserved across diverse and distantly related lineages of eukaryotes, with no closely related relatives to the taxa from this study. To expand this analysis further, we carried out our own manual analysis employing a similar approach. We annotated each MAG by training a specific Augustus model for each genome (see details below). Using these annotations and a set of ciliate proteomes (**Table S1**), we generated a set of ciliate orthogroups using OrthoFinder and identified in-frame UAG codons that correspond to highly conserved (≥ 70% identity) amino acid sites and recorded the most numerous amino acid at these sites. This analysis yielded the same predictions as PhyloFisher but with greater support using our ciliate-specific dataset. 1131, 1974, and 2431 in-frame UAG codons that correspond to highly conserved amino acid sites were identified in ARC_274, ARC_306, and SOC_66, respectively, of which 80.9%, 82.8%, and 87.7% corresponded to positions where leucine is highly conserved (**Fig. 1**).

**Figure 1.**
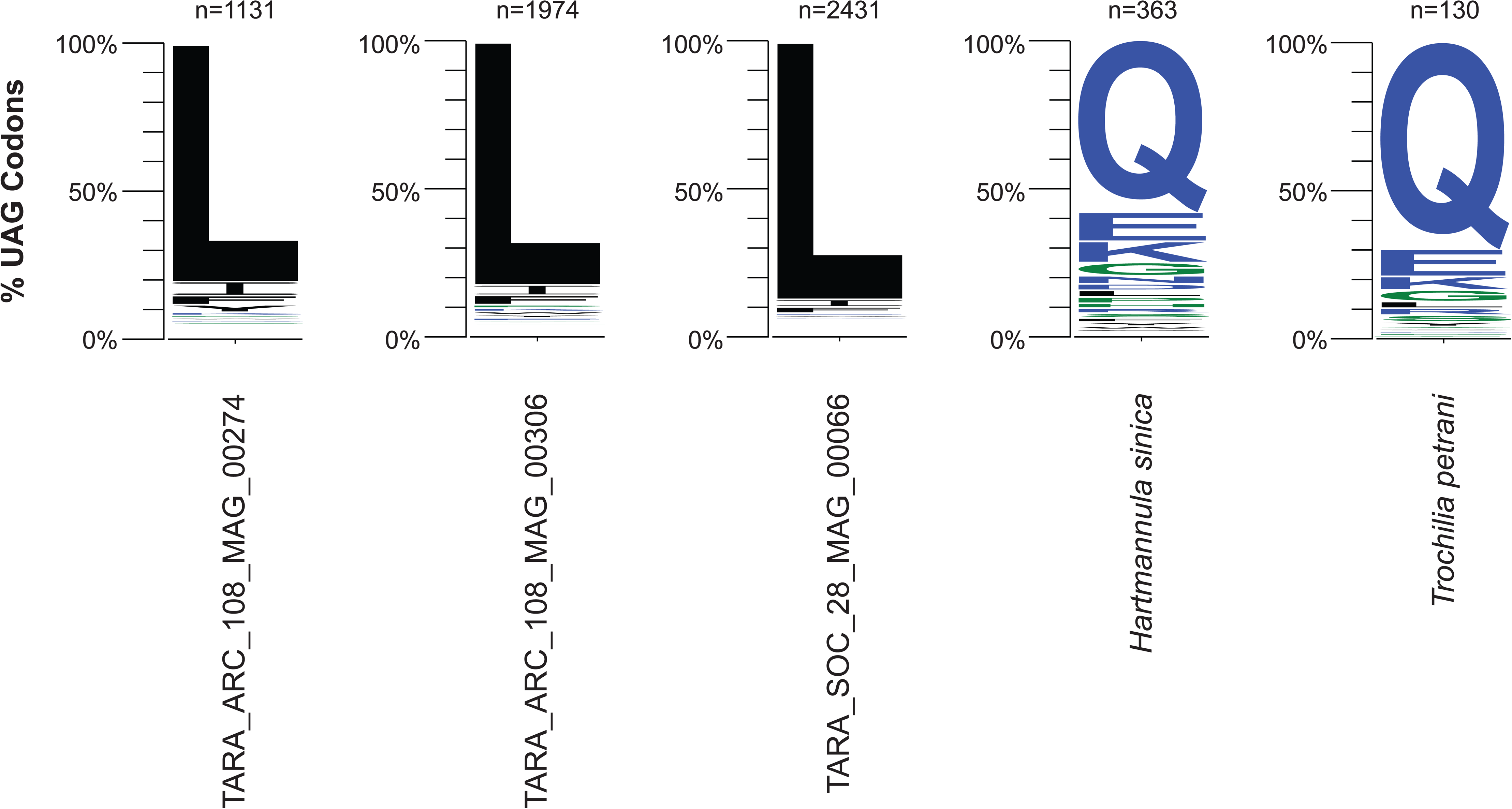
Genetic code prediction of the UAG codon. The genetic code of each species was predicted by identifying in-frame UAG codons that occur at highly conserved (≥ 70% identity) amino acid positions in ciliate orthogroups. Each sequence logo represents the frequency of the most numerous amino acids at these highly conserved positions for each species. Numbers represent the number of codons analysed.

An example multiple sequence alignment of the DRG2 protein (developmentally-regulated GTP-binding protein 2) is shown in **Fig 2** with ciliate sequences aligned against orthologs from diverse eukaryotic species. The ARC_274 sequence contains three internal UAG codons which correspond to positions where leucine is highly conserved in the other eukaryotic sequences (**Fig. 2**). The ARC_306 sequence contains a single internal UAG codon corresponding to leucine and the SOC_66 sequence contains three internal UAG codons corresponding to leucine (and one to isoleucine) (**Fig. 2**).

**Figure 2.**
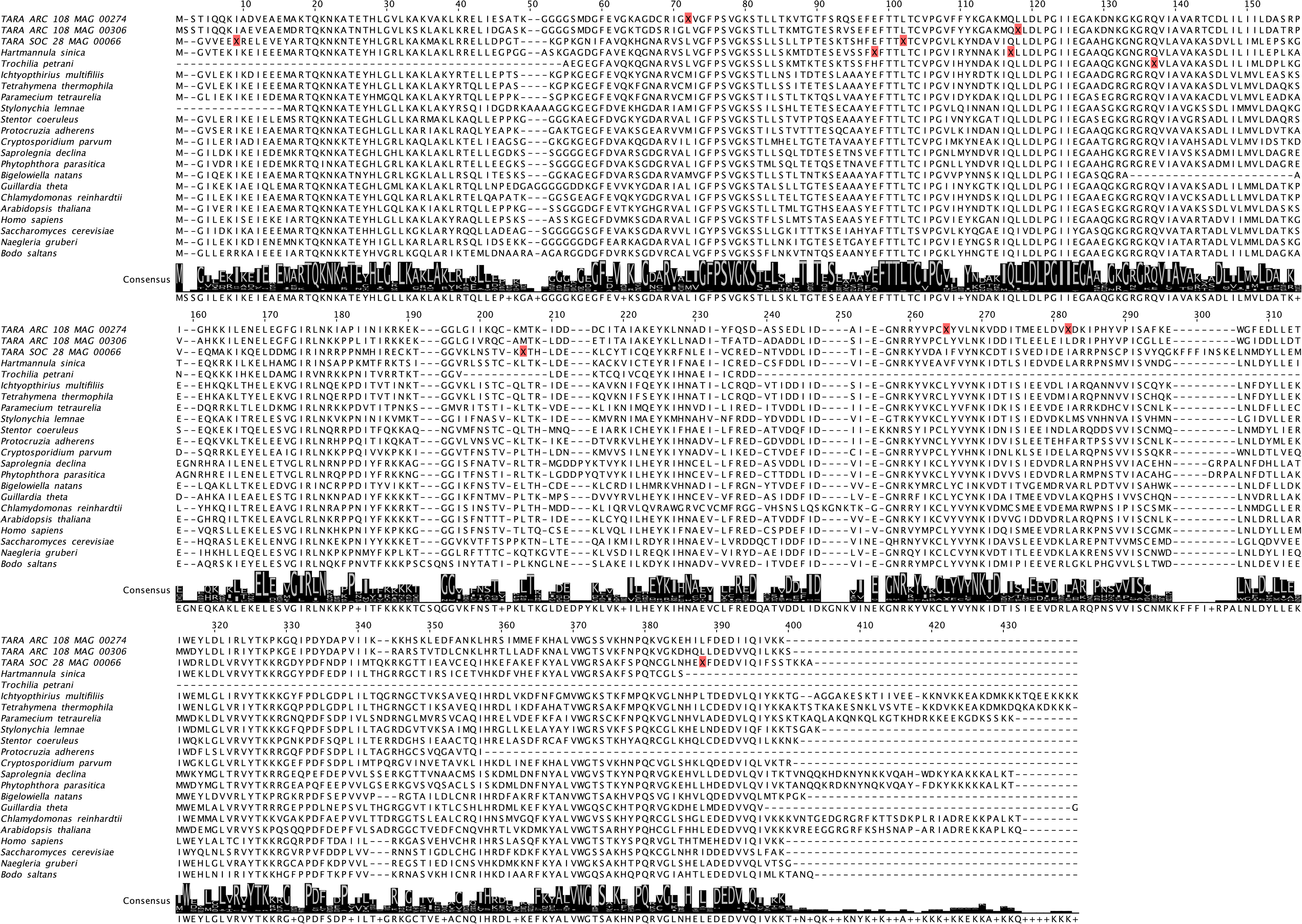
Example multiple sequence alignment of orthologs of the DRG2 protein (developmentally-regulated GTP-binding protein 2) from ciliates and diverse representatives across Eukaryota. Internal UAG codons are indicated by “X” with a red background. The in-frame UAG codons in the TARA MAGs occur at positions where leucine is highly conserved, whereas the in-frame UAG codons in *Hartmannula sinica* and *Trochilia petrani* occur at positions where glutamine is highly conserved.

A challenge with metagenome binning is that ribosomal rRNA genes are typically missing from MAGs, due to technical limitations, which is the case here making detailed taxonomic identification difficult. Instead, we relied upon the alpha-tubulin protein as a phylogenetic marker. An alpha-tubulin protein was recovered from just one of the MAGs (ARC_306). Phylogenetic analysis placed the ARC_306 sequence within a clade of sequences from the Phyllopharyngea class (**Fig. S3**) with high support (91% ultrafast bootstrap support). We extend this phylogenetic analysis below using phylogenomic approaches.

### Reassignment of the UAG stop codon to glutamine in *Hartmannula sinica* and *Trochilia petrani*

To expand our dataset further, we retrieved previously published genome sequencing reads from members of the Phyllopharyngea class (Maurer-Alcalá et al. 2018; Pan et al. 2019) and generated *de novo* genome assemblies for seven species – *Chilodochona* sp., *Chilodonella uncinata*, *Chilodontopsis depressa*, *Dysteria derouxi*, *Hartmannula sinica*, *Trithigmostoma cucullulus*, and *Trochilia petrani* (**Table S1**). We cleaned up each assembly to remove sequences from contaminants and predicted their genetic codes using the same methods as above. *Chilodochona* sp., *Chilodonella uncinata*, *Chilodontopsis depressa*, *Dysteria derouxi*, and *Trithigmostoma cucullulus* were all predicted use the canonical genetic code with both methods (**Fig. S1**). Surprisingly, however, we predicted that *Hartmannula sinica* and *Trochilia petrani* use non-canonical genetic codes, where the UAG stop codon has been reassigned to code for glutamine. Codetta identified 1489 and 350 UAG codons with a Pfam alignment in *Hartmannula sinica* and *Trochilia petrani*, respectively, and predicted that they translate UAG to glutamine with a log decoding probability of zero (**Fig. S1**). This prediction was supported by PhyloFisher which identified 30 and 7 internal UAG codons in *Hartmannula sinica* and *Trochilia petrani*, respectively, which correspond to positions where glutamine is highly conserved (**Fig. S2**). Furthermore, our manual analysis of predicted gene models and ciliate orthogroups identified 363 and 130 internal UAG codons in *Hartmannula sinica* and *Trochilia petrani*, respectively, of which 58.1% and 70.0% correspond to positions where glutamine is highly conserved (**Fig. 1**).

The partial *Trochilia petrani* DRG2 sequence contains a single internal UAG codon corresponding to glutamine and the *Hartmannula sinica* DRG2 sequence contains a single internal UAG codon corresponding to glutamine (and one to glutamate) (**Fig. 2**). The five species with the canonical genetic code had high BUSCO completeness (72.5% to 82.5%) but the two species with reassigned UAG codons had low completeness (30.4% for *Hartmannula sinica* and 8.8% for *Trochilia petrani*) (**Table S1**).

### Phylogenomics reveals three independent genetic codon reassignments in Phyllopharyngea

To better understand the evolutionary relationships of the three uncultivated TARA MAGs, and to characterise the order of events surrounding the novel genetic code reassignments, we carried out a phylogenomics analysis focused on members of the CONthreeP lineage of ciliates – Colpodea, Oligohymenophorea, Nassophorea, Phyllopharyngea, Plagiopylea, and Prostomatea (which isn’t represented here). A concatenated alignment of 115 BUSCO proteins (53,648 amino acid sites after alignment trimming) from 29 ciliate species was constructed and used for phylogenomic analyses. Phylogenomic reconstruction was performed using maximum-likelihood (ML) and Bayesian approaches. The ML analysis was conducted using IQ-Tree under the LG+C20+F+G+PMSF model with 100 non-parametric bootstraps, while the Bayesian analysis was conducted using PhyloBayes MPI under the CAT-GTR model. Both methods yielded robust phylogenies with identical topologies and all branches had full statistical support from both methods (i.e., ML bootstrap support of 100% and a Bayesian posterior probability of 1) (**Fig. 3**). Our phylogeny is in agreement with previous phylogenies based on small subunit ribosomal rRNA genes (Gao et al. 2012; Pan et al. 2019).

**Figure 3.**
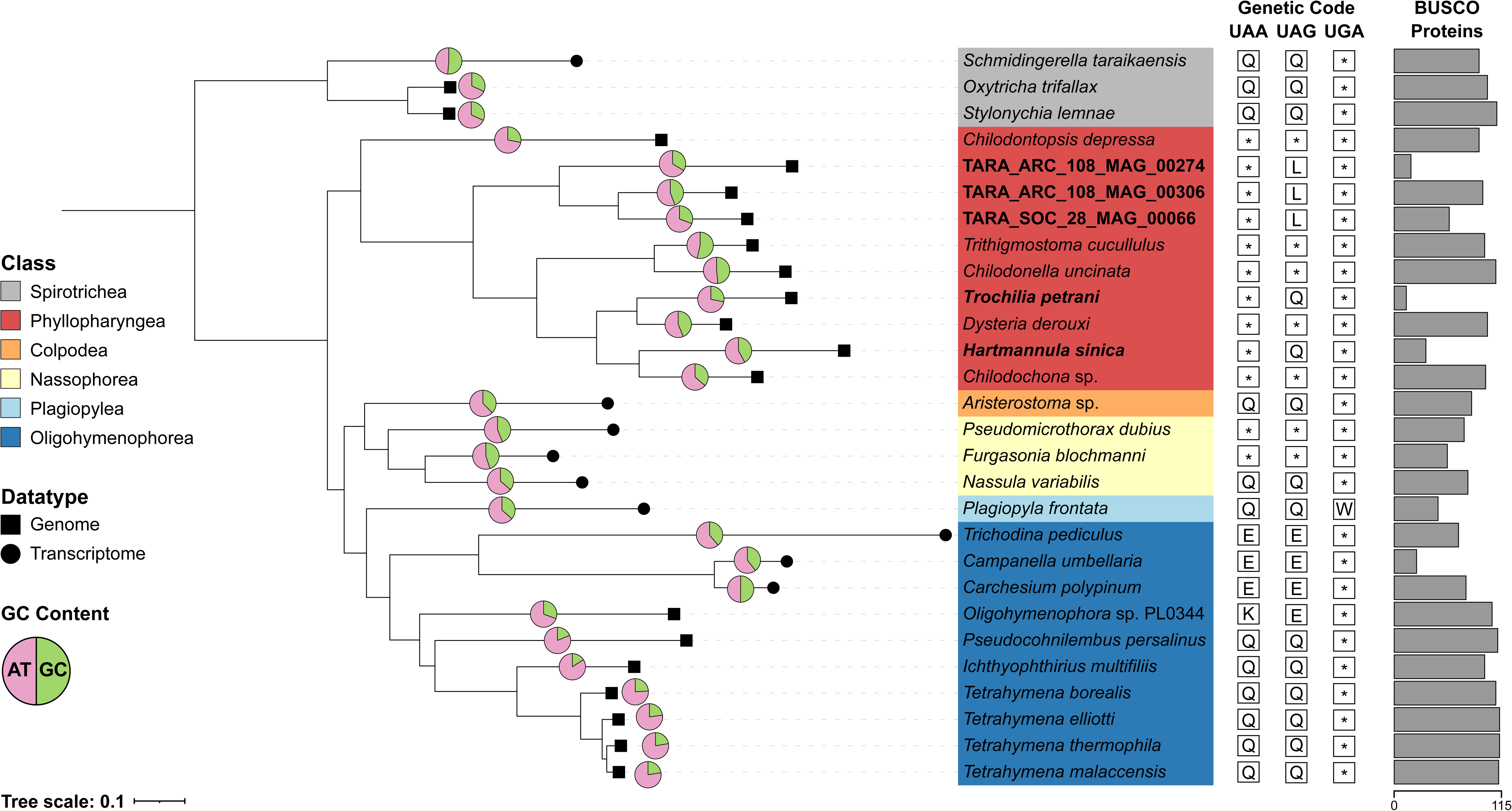
Phylogenomics analysis of the CONthreeP lineage of ciliates. The phylogeny was reconstructed from a concatenated alignment of 115 BUSCO proteins (53,648 amino acid sites). Maximum-likelihood analysis was conducted using IQ-Tree under the LG+C20+F+G model with PMSF approximation and 100 non-parametric bootstraps. Bayesian inference was performed using PhyloBayes MPI using the CAT-GTR model. The branch lengths displayed are from the ML analysis. All branches have full statistical support from both methods (i.e., ML bootstrap support of 100% and a Bayesian posterior probability of 1). *Oxytricha trifallax*, *Schmidingerella taraikaensis*, and *Stylonychia lemnae* are included as outgroups from the Spirotrichea class. The type of data (genomic or transcriptomic) is indicated by symbols at branch tips. GC content for each species is shown in a pie chart. Note that caution is required when comparing GC content between genome and transcriptome assemblies. The number of BUSCO proteins included in the concatenated alignment is shown in the bar pot, highlighting the amount of missing data per species. Genetic code changes are shown (*, STOP; Q, glutamine; L, leucine; K, lysine; W, tryptophan; E, glutamic acid).

The three TARA MAGs formed a monophyletic lineage within Phyllopharyngea (**Fig. 3**), confirming that they belong to the Phyllopharyngea class. This suggests that the reassignment of UAG to leucine occurred once in an ancestor of this lineage. ARC_306 grouped as sister to SOC_66, to the exclusion of ARC_274, despite geographical differences. *Trochilia petrani* and *Dysteria derouxi* were grouped as sister lineages, as were *Hartmannula sinica* and *Chilodochona* sp. **(Fig. 3**). This suggests that translation of UAG to glutamine independently evolved twice within sampled Phyllopharyngea species and that these genetic code changes were more recent than the reassignment of UAG to leucine in the TARA MAGs lineage. The phylogenetic distribution of species that use the canonical genetic code suggests that the most recent common ancestor of Phyllopharyngea used the canonical genetic code (**Fig. 3**).

### Novel suppressor tRNAs for UAG

Translation of the UAG codon to an amino acid requires a tRNA gene that can decode the UAG codon. We annotated tRNA genes in our dataset using tRNAscan-SE (Chan et al. 2021). A single suppressor tRNA gene was identified in the ARC_274 MAG with a CUA anticodon which is predicted with high confidence to function as a leucine tRNA (**Fig. S4A**). This tRNA gene is located on a 12.5 kb contig, capped at both ends with ciliate telomeric repeats (CCCCAAA/GGGGTTT). Two suppressor tRNA genes were identified in the ARC_306 MAG with CUA anticodons that were predicted with high confidence to function as leucine tRNAs (**Fig. S4A**). Both of these suppressor tRNA genes are located on the same 13.5 kb contig within 100 bp of each other but are not identical (75% identical) (**Fig. S4B**). This contig has three gene models with best BLAST hits to ciliate sequences. One of the ARC_306 suppressor tRNAs is 94% identical to the suppressor tRNA from ARC_274, with only five nucleotide differences between the two sequences (one substitution in the anticodon loop and four substitutions in the variable loop) (**Fig. S4A, Fig. S4B**). We compared the suppressor tRNA gene sequences against other phyllopharyngean tRNA sequences which revealed that they are most similar to leucine tRNAs with CAA or TAA anticodons (**Fig. S4B**). This suggests that the novel suppressor tRNAs evolved from a canonical leucine tRNA. We did not detect suppressor tRNA genes in the SOC_66 MAG, *Hartmannula sinica*, or *Trochilia petrani,* however our analysis is likely limited by the low completeness of these assemblies. As expected, suppressor tRNA genes were not detected in the five genomes that use the standard genetic code.

### Codon usage of canonical and reassigned stop codons

To better understand the events preceding a genetic code change event, we annotated all 10 phyllopharyngean genomes and investigated usage of the canonical stop codons and the reassigned UAG codon following a genetic code change. We restricted this analysis to exclude partial gene models by only considering genes with both a predicted start and stop codon. UAA is the most used stop codon and UGA is the least used stop codon in all species in our dataset (**Fig. 4**). *Trithigmostoma cucullulus* is a clear outlier in terms of codon usage with 58.8% of genes using UAA, 30.3% using UAG, and 10.9% using UGA as stop codons (**Fig. 4**), which is reflected by the higher GC content of its genome (**Fig. 3**) (**Table S1**). UAA is used as stop codon in 78.3% – 86% of genes in the other species that use the standard genetic code (**Fig. 4**). In the species that have retained UAG as a stop codon (excluding *Trithigmostoma*), UAG is used as stop codons for 10.5% – 16.2% of genes (**Fig. 4**). This suggests that UAG usage was low but non-negligible prior to the genetic code change events. UGA is used less frequently in these species, with only 3.1% – 9.4% of genes using UGA as a stop codon (**Fig. 4**).

**Figure 4.**
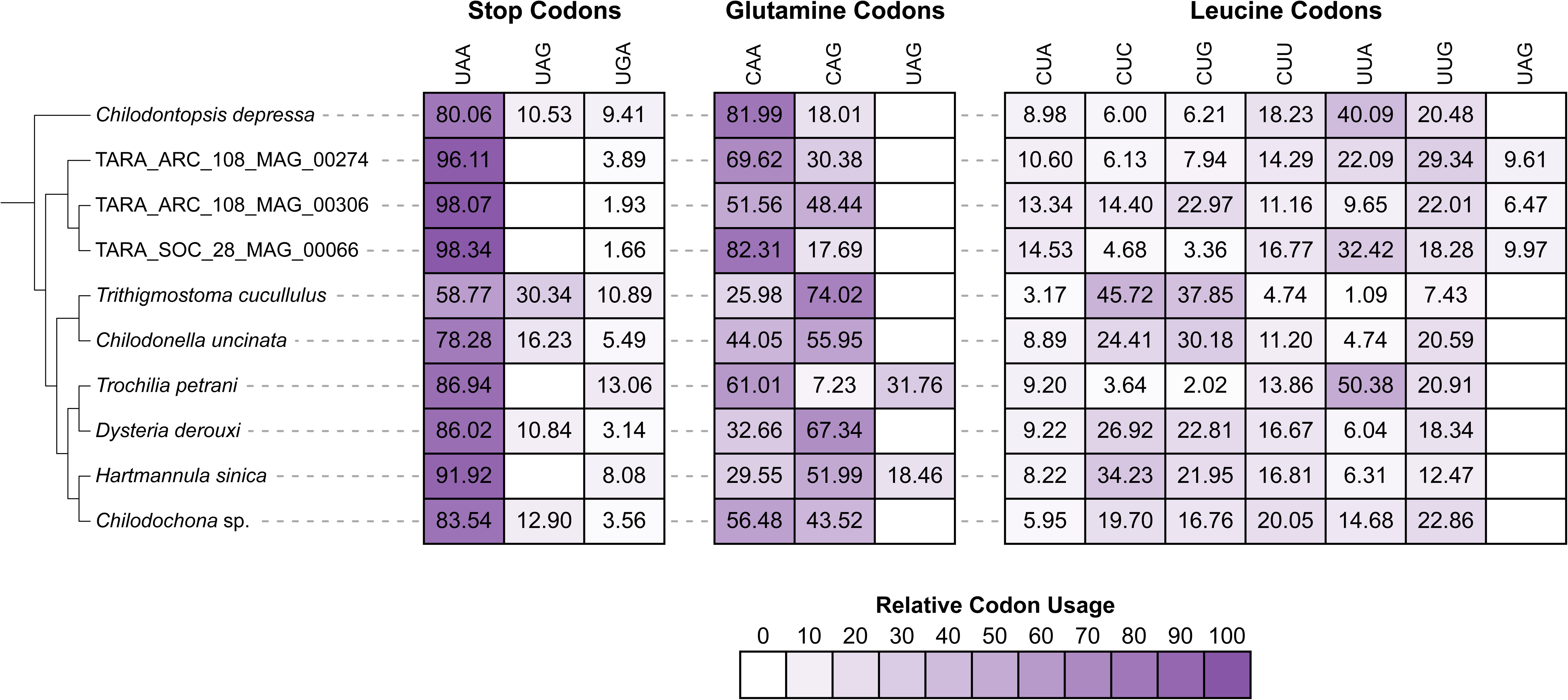
Codon usage of canonical stop, glutamine, and leucine codons, and the reassigned UAG codons. The heatmap shows the relative usage of each codon. The cladogram depicts the phylogenetic relationships determined in our phylogenomic analysis in Fig. 3.

Usage of UAA as a stop codon is increased to 96.1% – 98.3% of genes in the TARA MAGs following their genetic code reassignment (**Fig. 4**). Fewer than 3.9% of genes use UGA as a stop codon in these MAGs. UAA usage is also increased in *Hartmannula sinica* compared to its closest relative (91.9% vs 83.5%) (**Fig. 4**). Likewise, UGA usage increased in *Hartmannula sinica* (8.1% vs 3.6%) and *Trochilia petrani* compared to their closest relatives (13.1% vs 3.1%) following their genetic code reassignments (**Fig. 4**).

In *Trochilia petrani* and *Hartmannula sinica*, we compared relative codon usage of the reassigned UAG codon compared to the two canonical glutamine codons (CAA and CAG). The reassigned UAG codon is the second most used glutamine codon in *Trochila petrani* (31.8%) but the least used in *Hartmannula sinica* (18.5%) (**Fig. 4**). 83.7% and 78.2% of genes in *Trochila* and *Hartmannula* contain at least one internal in-frame UAG codon, showing that the reassigned codon is widely used in both species. In the three TARA MAGs, we compared usage of the reassigned UAG codon compared to the six canonical leucine codons (CUA, CUC, CUG, CUU, UUA, and UUG). Relative codon usage of the reassigned UAG codon ranges from 6.5% to 10% in the TARA MAGs (**Fig. 4**). 81.3% to 91.1% of genes contain at least one internal UAG codon in these MAGs, showing that usage of the reassigned codon is also widespread in these genomes.

## Discussion

Here we report the identification of three novel genetic code changes in ciliates. We discovered that three uncultivated ciliates sequenced from eukaryotic metagenomes by the TARA Oceans Project (Delmont et al. 2022) use a non-canonical genetic code where the UAG stop codon has been reassigned to encode leucine (**Fig. 1**). Phylogenomic analysis revealed that the uncultivated ciliates belong to the Phyllopharyngea class. Reassembly and analysis of seven other phyllopharyngean genomes led to the discovery of another two genetic code changes in this lineage – reassignment of UAG to glutamine in both *Hartmannula sinica* and *Trochilia petrani* (**Fig. 1**).

It is important to note that the genetic code changes reported herein, and indeed most published reports of genetic code changes, are predictions based on genomic data. While these are high-confidence predictions with multiple lines of evidence including large numbers of internal in-frame UAG codons that occur at conserved leucine/glutamine positions and the identification of predicted cognate suppressor tRNA genes in two genome assemblies, confirmation that a codon is translated to a particular amino acid requires proteome sequencing (e.g., mass spectrometry) and comparison with corresponding coding sequences.

Given the complex phylogenetic distribution of canonical and non-canonical genetic codes in the ciliate lineage (McGowan et al. 2023), there are conflicting interpretations surrounding the order of events. Either (1) independent genetic code changes occurred in multiple ciliate lineages, including different lineages convergently evolving the same non-canonical genetic code variant, or (2) stop codon reassignments evolved in more ancient lineages of ciliates giving rise to the taxa with non-canonical genetic codes, which were followed by reversions to the original function as stop codons in the taxa that use the standard genetic code. A recent study proposed that ancestral ciliates reassigned all three stop codons as sense codons, followed by one or more of the reassigned codons reverting to functioning as stop codons giving rise to the different types of genetic codes observed in ciliates (i.e., the standard genetic code or genetic codes with one or more reassigned stop codons) (Chen et al. 2023). In our study, we focus just on the Phyllopharyngea class of ciliates. From these phylogenomic analyses, the most parsimonious explanation for the distribution of genetic codes within sampled phyllopharyngean ciliates is that the most recent common ancestor of Phyllopharyngea used the canonical genetic code (**Fig. 3**). This lineage then underwent at least three independent genetic code change events based on sampled species: (1) reassignment of UAG to leucine in the lineage of uncultivated TARA MAGs, (2) reassignment of UAG to glutamine in the lineage giving rise to *Hartmannula sinica*, and (3) reassignment of UAG to glutamine in the lineage giving rise to *Trochilia petrani* (**Fig. 3**). Given the widespread usage of the reassigned UAG codons in Phyllopharyngea, with 78.2% to 91.1% of genes using a least one UAG codon, it is unlikely that UAG will revert to functioning as a stop codon. If it were to revert, it would result in large-scale protein truncation due to the presence of internal in-frame UAG codons in most genes unless there were genome-wide substitutions of UAG to another sense codon (synonymous or non-synonymous). Thus, we propose that it is unlikely that a reversion could so readily occur once a stop codon has been reassigned to a sense codon.

Several hypotheses have been proposed to model the processes surrounding genetic code changes. Under the “codon capture” hypothesis, a codon is driven to extinction by mutational biases (e.g., low GC-content) followed by loss of the corresponding tRNA (or loss of function in release factors in the case of stop codons) (Osawa and Jukes 1989). This unused codon could later be captured by a noncognate tRNA and reappear in the genome, resulting in a change to the genetic code. The “ambiguous intermediate” hypothesis proposes that genetic code changes involve an intermediate stage where a codon is ambiguously translated via competing tRNAs charged with different amino acids (Schultz and Yarus 1994). In the scenario of a stop codon being reassigned, this would involve a suppressor tRNA competing with a release factor protein. The “tRNA loss driven codon reassignment” hypothesis proposes that the genetic code change is preceded by loss of function or reduced efficiency of a tRNA or release factor, resulting in an unassigned or inefficient codon that can be captured by another tRNA gene (Mühlhausen et al. 2016).

It is still unclear what has driven ciliates to evolve so many genetic code variants and why there is differential retention of UAA/UAG/UGA as stop codons. Our analysis of codon usage shows that usage of UAG as a stop codon is low but non-negligible (mean 16.2% of genes) in phyllopharyngean species that use the canonical genetic code (**Fig. 4**). UGA is the least used stop codon in all taxa in our dataset (1.7% - 13.1% of genes) (**Fig. 4**). Thus, it is unclear why UGA was retained as a stop codon but not UAG. Furthermore, the UGA codon is known to be the least robust and most prone to translational readthrough (Dabrowski et al. 2015) confounding the situation further. Likewise, the low GC content that is a common characteristic of extant ciliate genomes does not explain their propensity to evolve non-standard genetic codes. GC content varies considerably within Phyllopharyngea (27.4% to 53.6%) (**Fig. 3**). *Chilodontopsis depressa* has a lower GC content (28%) than most of the other species in our dataset but uses the standard genetic code and has similar stop codon usage as other species with higher GC content (**Fig. 3 and 4**).

The evolution of the UAA and UAG codons are thought to be coupled as they virtually always have the same meaning – either they both function as canonical stop codons or they are both reassigned to code for the same amino acid (Kollmar and Mühlhausen 2017). The first reported cases where UAA and UAG have different meanings in a nuclear genome were reported in two unrelated taxa – an uncultured Rhizarian where UAG was reassigned to code for leucine and in the Fornicate *Iotanema spirale* where UAG was recoded for glutamine (Pánek et al. 2017). Recently, we reported a novel genetic code variant in an uncultured ciliate, where the UAA and UAG codons were reassigned to code for two different amino acids (lysine and glutamic acid, respectively), the first reported case where UAA and UAG encode two different amino acids (McGowan et al. 2023). In this study, we report another deviation from the trend of UAA and UAG having the same function. We show that UAG has been reassigned to function as a sense codon, but UAA was retained as a stop codon in five of the studied genomes. Furthermore, we show that it is unlikely that the UAA stop codon could later be reassigned to an amino acid in these lineages, as seen in most other non-canonical genetic code variants, given the almost ubiquitous usage of UAA as the preferred stop codon with 86.9% to 98.3% of genes using UAA as a stop codon (**Fig. 4**). If UAA were to be later reassigned to function as a sense codon, it would result in widespread protein elongation with proteins extending downstream to the next in-frame stop codon (i.e., UGA). Ciliates typically have very short 3’-UTRs which would somewhat limit the effect, but this would still impact almost the entire proteome. Such levels of protein elongation would likely have similar deleterious consequences as widespread translational readthrough, including issues with protein aggregation and stability, disruption to localisation signals and energetic waste (Ho and Hurst 2022). Thus, the function of UAA as the preferred stop codon is likely fixed in these taxa. Had the UAA codon been reassigned initially, it likely would have also triggered the UAG codon to be reassigned to encode the same amino acid given that a suppressor tRNA that decodes UAA is also expected to decode UAG due to wobble base pairing of the UUA anticodon (Hanyu et al. 1986). This is not the case when UAG is reassigned on its own, as a suppressor tRNA with an anticodon complementary to UAG (i.e., CUA anticodon) is not expected to recognise the UAA codon, allowing the UAG codon to evolve independently of UAA.

Our results highlight the evolvability of the genetic code and the tendency, not only for ciliates but also unrelated taxa, to independently evolve the same non-canonical genetic code variants. Multiple lineages within Ciliophora have independently evolved the translation of UAR codons to glutamine (McGowan et al. 2023). Translation of UAR to glutamine is also found in the nuclear genomes of green algae from the Ulvophyceae class (Cocquyt et al. 2010), the diplomonad *Hexamita* (Keeling and Doolittle 1996), the oxymonad *Streblomastix strix (Keeling and Leander 2003)*, and the aphelid *Amoeboaphelidium protococcarum* (Karpov et *al. 2013*). Similarly, reassignment of the UAG stop codon (but not UAA) to leucine is reported here for the first time in ciliates but was previously reported in the nuclear genome of an uncultured Rhizarian (Pánek et al. 2017) and also in the mitochondrial genomes of chlorophyte algae (Hayashi-Ishimaru et al. 1996; Noutahi et al. 2019) and in some chytridiomycete fungi (Laforest et al. 1997). Reassignment of UAG (but not UAA) to glutamine is also reported here to have evolved independently twice within sampled phyllopharyngean ciliates and was also previously reported in the Fornicate *Iotanema spirale* (Pánek et al. 2017). These findings highlight that while the genetic code is one of the most conserved features of molecular biology, it is not quite as universal as was once thought (Hinegardner and Engelberg 1963). Non-canonical genetic code reassignments are relatively recent events, demonstrating that genetic code evolution is an ongoing process.

## Materials and methods

### Dataset assembly

Ciliate MAGs from the TARA Oceans project were downloaded from https://www.genoscope.cns.fr/tara/ (Delmont et al. 2022). The three MAGs focused on in this study are TARA_ARC_108_MAG_00274, TARA_ARC_108_MAG_00306, and TARA_SOC_28_MAG_00066. Genome sequencing reads for *Chilodochona* sp. (SRR9841583), *Chilodontopsis depressa* (SRR9841577), *Dysteria derouxi* (SRR9841578), *Hartmannula sinica* (SRR9841582), *Trithigmostoma cucullulus* (SRR9841579), and *Trochilia petrani* (SRR9841580) were downloaded from BioProject PRJNA546036 (Pan et al. 2019). Genome sequencing reads for *Chilodonella uncinata* (SRR6195035) were downloaded from BioProject PRJNA413041 (Maurer-Alcalá et al. 2018).

Genome assemblies were generated using SPAdes (v3.15.5) (Prjibelski et al. 2020) with default settings, except single-cell (--sc) mode was enabled. Contigs shorter than 1000 bp were discarded. Assemblies were decontaminated using a combination of Tiara (Karlicki et al. 2022) and contig clustering based on tetranucleotide frequencies.

### Genetic code prediction and genome annotation

The genetic code used by each genome was initially predicted using Codetta (v2.0) (Shulgina and Eddy 2023) and the PhyloFisher “genetic_code_examiner” utility (Tice et al. 2021). For the five phyllopharyngean species that use the canonical genetic code, an initial gene set was generated using GeneMark-EP (Brůna et al. 2020), with hints generated by ProtHint from a database of 1,170,806 Alveolata proteins. These initial gene sets were filtered by selecting complete gene models (i.e., containing a start and stop codon) that had full-length alignments (alignment length ≥ 95% of both the query and subject sequence lengths) against the Alveolata protein database using Diamond in ultra-sensitive mode (Buchfink et al. 2021). These filtered subsets were used as training gene sets to train an Augustus model for each species (Stanke et al. 2006). This process was repeated by incorporating the initial Augustus gene models from every other species into the protein database supplied to GeneMark-EP and to retrain Augustus and generate a final gene set for each species. For the five species that use a non-canonical genetic code, an initial gene set was generated using the most appropriate Augustus model from above (with modified parameters such that UAG is no longer used as a stop codon), which went through a similar filtering step to select training genes to train a species-specific Augustus model for each genome.

The final gene sets were used to further interrogate the genetic code. In-frame UAG codons were translated to “X”. Orthogroups from a dataset of 19 ciliate species (including the 10 phyllopharyngean genomes) (**Table S1**) were identified using OrthoFinder (v.2.5.5) (Emms and Kelly 2019) with the parameter “-M msa”. A multiple sequence alignment was generated for each orthogroup using MAFFT (Katoh and Standley 2013). In-frame UAG codons that correspond to highly conserved amino acid positions (≥ 70% identity) in aligned orthogroups were identified and the most numerous amino acid at these sites were counted. The counts were visualised as a sequence logo using WebLogo (v3.7.12) (Crooks et al. 2004). This genetic code analysis and subsequent analyses of codon usage were restricted to gene models with predicted start and stop codons (i.e., not partial gene models). tRNA genes were annotated using tRNAscan-SE (Chan et al. 2021).

### Phylogenetics of alpha-tubulin sequences

An alpha-tubulin gene was recovered from ARC_306 and used for phylogenetic analysis, using a dataset of alpha-tubulin sequences from a previously published phylogenetics study of ciliates (Gao et al. 2016). Three apicomplexan sequences were included as outgroups. Sequences were aligned using MAFFT (v7.520) with the L-INS-I algorithm (Katoh and Standley 2013). A maximum-likelihood phylogeny was constructed using IQ-Tree (v2.2.2.6) (Minh et al. 2020) under the LG+R3 model which was the best fitting model according to ModelFinder (Kalyaanamoorthy et al. 2017). Support was assessed using 1000 ultrafast bootstrap replicates (Hoang et al. 2018).

### Phylogenomics

We constructed a dataset of ciliate genomes and transcriptomes for phylogenomic analyses, focusing on members of the CONthreeP lineage (**Fig. 3**). *Oxytricha trifallax*, *Schmidingerella taraikaensis*, and *Stylonychia lemnae* from the Spirotrichea class were included as outgroups. A concatenated supermatrix of BUSCO proteins was generated using our BUSCO_phylogenomics pipeline (https://github.com/jamiemcg/BUSCO_phylogenomics). 115 BUSCO proteins from the Alveolata_odb10 dataset (Manni et al. 2021) were identified as complete and single copy in at least 60% of species and were included in our analysis. Each BUSCO family was individually aligned using MUSCLE (v5.1) (Edgar 2022). Alignments were then trimmed using trimAl (v1.4) (Capella-Gutierrez et al. 2009) with the “automated1” parameter and concatenated together resulting in a supermatrix alignment of 53,648 sites. Maximum-likelihood phylogenomic reconstruction was performed using IQ-TREE (v2.2.2.6) (Minh et al. 2020) under the LG+C20+F+G model with PMSF approximation (Wang et al. 2018) and 100 non-parametric bootstraps, using a guide tree from FastTree (v2.1.11) (Price et al. 2010). Bayesian analyses were also conducted using PhyloBayes MPI (v1.8) (Lartillot et al. 2013) under the CAT-GTR model. Two independent Markov chain Monte Carlo chains were run for approximately 10,000 generations. Convergence was assessed using bpcomp and tracecomp, with a burn-in of 20%. Phylogenies were visualised and annotated using iTOL (Letunic and Bork 2021).

## Acknowledgements

This work was funded by the Wellcome Trust Darwin Tree of Life Awards (218328 and 226458) and by the Biotechnology and Biological Sciences Research Council (BBSRC), part of UK Research and Innovation, through the Core Capability Grant (BB/CCG2220/1) at the Earlham Institute; the Earlham Institute Strategic Programme Grant Decoding Biodiversity (BBX011089/1) and its constituent work packages (BBS/E/ER/230002A and BBS/E/ER/230002B). The authors acknowledge the work delivered via the Research Computing Group at EI who manage and deliver High Performance Computing at EI. TAR is supported by a Royal Society University Research Fellowship (URF/R/191005).

## Data Availability

Supporting data have been deposited on Zenodo (10.5281/zenodo.12744466).

---

**Figure S1.** Genetic code prediction of the UAG codon using Codetta. The table shows log decoding probabilities of UAG for each amino acid. “?” indicates that there were insufficient alignments to infer an amino acid meaning (which is the expected behaviour for stop codons).

**Figure S2.** Genetic code prediction of the UAG codon using the genetic_code_examiner utility from PhyloFisher.

**Figure S3.** Maximum-likelihood phylogeny of 104 ciliate alpha-tubulin sequences constructed using IQ-TREE under the LG+R3 model. Numbers represent support from 1000 ultrafast bootstrap replicates. Three apicomplexan sequences were included as an outgroup.

**Figure S4. (A)** Three suppressor tRNA genes from two TARA Oceans ciliate MAGs (ARC_274 and ARC_306) with CUA anticodons that are predicted to function as leucine tRNAs. **(B)** Multiple sequence alignment of the three suppressor tRNA genes with representative canonical leucine tRNAs with CAA and TAA anticodons.

## Table Captions

**Table S1.** Summary of the genome assembly and annotation statistics and the species included in the OrthoFinder analysis.

