## Supplemental Figure 1 for "Multiple Independent Genetic Code Reassignments of the UAG Stop Codon in Phyllopharyngean Ciliates"

| | A | C | D | E | F | G | H | I | K | L | M | N | P | Q | R | S | T | V | W | Y | ? |
|---|---|---|---|---|---|---|---|---|---|---|---|---|---|---|---|---|---|---|---|---|---|
| *Chilodontopsis depressa* | -40.81 | -85.74 | -30.44 | -28.35 | -72.81 | -56.66 | -39.97 | -65.48 | -33.56 | -62.16 | -60.60 | -27.23 | -70.06 | -18.73 | -46.94 | -27.77 | -33.36 | -63.66 | -101.55 | -62.09 | **0.00** |
| TARA_ARC_108_MAG_00274 | -2091.97 | -2351.75 | -4215.32 | -3690.67 | -1488.93 | -3845.54 | -3169.67 | -697.02 | -3394.18 | **0.00** | -826.24 | -3409.71 | -3578.15 | -2949.29 | -3220.11 | -2813.36 | -2109.52 | -979.00 | -3050.52 | -2468.75 | -545.60 |
| TARA_ARC_108_MAG_00306 | -3871.08 | -4700.99 | -7843.32 | -6851.27 | -2876.66 | -7224.96 | -5897.50 | -1535.69 | -6282.83 | **0.00** | -1680.20 | -6410.23 | -6695.74 | -5380.27 | -6097.99 | -5253.27 | -3972.26 | -2016.61 | -5948.80 | -4664.07 | -952.10 |
| TARA_SOC_28_MAG_00066 | -5742.39 | -6533.12 | -11407.97 | -10103.23 | -4164.23 | -10155.00 | -8535.61 | -2221.70 | -9368.85 | **0.00** | -2421.30 | -9430.68 | -9733.98 | -7911.45 | -8972.76 | -7766.88 | -5986.85 | -2940.86 | -8592.77 | -6926.17 | -1648.31 |
| *Trithigmostoma cucullulus* | -25.62 | -53.91 | -29.53 | -22.88 | -52.83 | -31.17 | -26.00 | -33.58 | -17.40 | -40.01 | -41.29 | -20.16 | -45.94 | -19.37 | -24.64 | -17.52 | -19.05 | -28.76 | -60.93 | -44.98 | **0.00** |
| *Chilodonella uncinata* | -27.75 | -55.64 | -45.10 | -30.26 | -36.38 | -68.98 | -32.25 | -48.25 | -36.52 | -40.96 | -34.32 | -25.47 | -58.73 | -22.31 | -36.99 | -22.61 | -26.07 | -38.80 | -55.16 | -27.98 | **0.00** |
| *Trochilia petrani* | -627.48 | -1223.45 | -699.93 | -443.11 | -1208.84 | -911.80 | -572.02 | -1158.34 | -474.31 | -1008.62 | -883.22 | -516.56 | -1097.49 | **0.00** | -569.01 | -498.23 | -584.84 | -946.08 | -1419.01 | -969.87 | -264.44 |
| *Dysteria derouxi* | -27.69 | -86.14 | -26.97 | -7.20 | -91.22 | -63.46 | -18.01 | -68.84 | -10.89 | -59.47 | -52.23 | -16.74 | -57.81 | -5.37 | -15.83 | -18.63 | -24.38 | -42.87 | -93.35 | -65.90 | **-0.01** |
| *Hartmannula sinica* | -2143.62 | -4569.62 | -2454.62 | -1438.33 | -4567.37 | -3440.24 | -2213.33 | -4047.66 | -1636.42 | -3509.28 | -3109.71 | -1895.60 | -3828.68 | **0.00** | -1904.59 | -1845.04 | -2088.89 | -3468.06 | -5193.11 | -3809.87 | -933.01 |
| *Chilodochona* sp | -16.11 | -53.56 | -42.09 | -26.09 | -48.72 | -42.03 | -23.36 | -30.42 | -14.44 | -33.17 | -35.05 | -19.50 | -43.18 | -23.25 | -25.05 | -11.40 | -8.59 | -22.90 | -80.15 | -46.11 | **0.00** |
