## Supplementary figures and images for "Multiple Independent Genetic Code Reassignments of the UAG Stop Codon in Phyllopharyngean Ciliates"

### Supplemental Figure 2

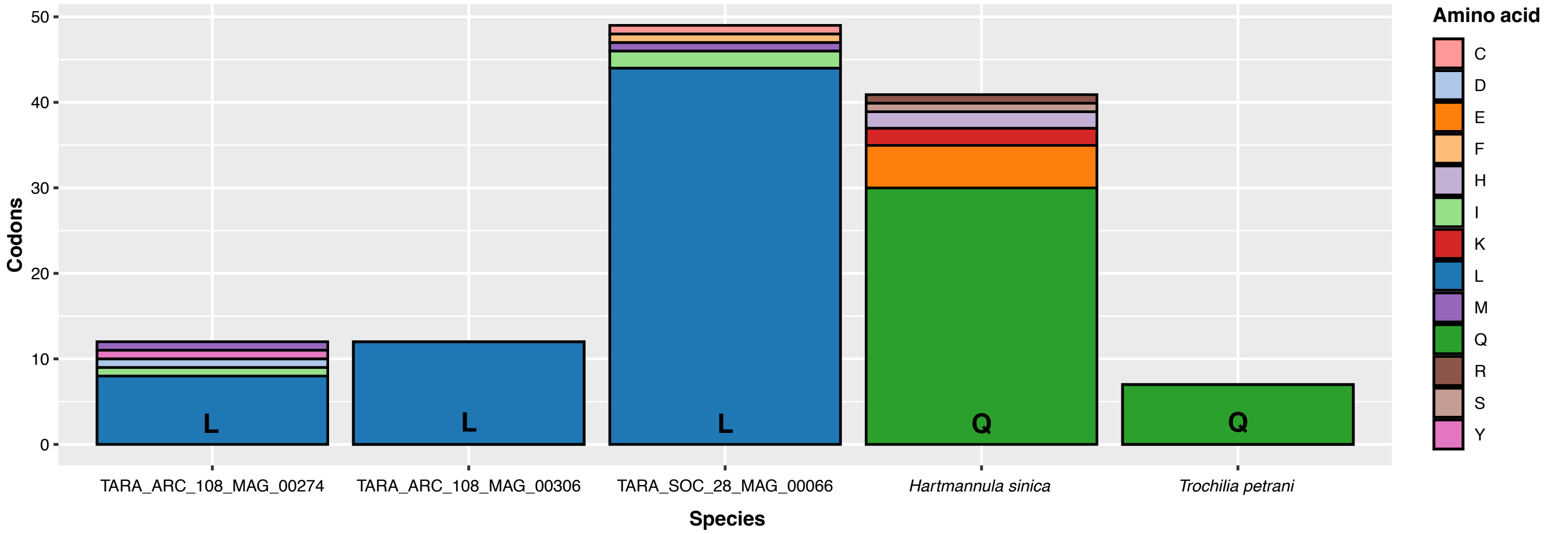

### Supplemental Figure 3

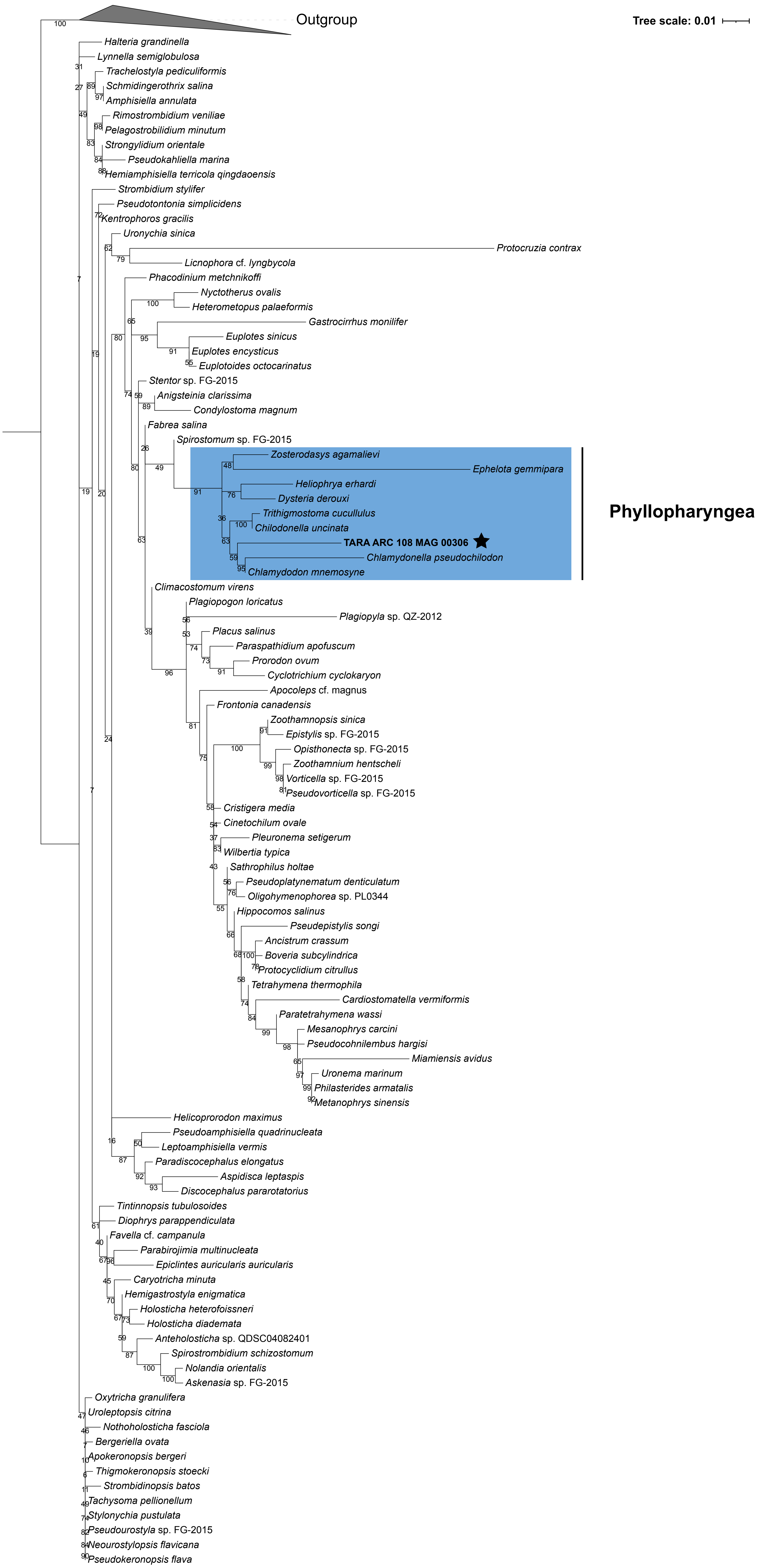

### Supplemental Figure 4

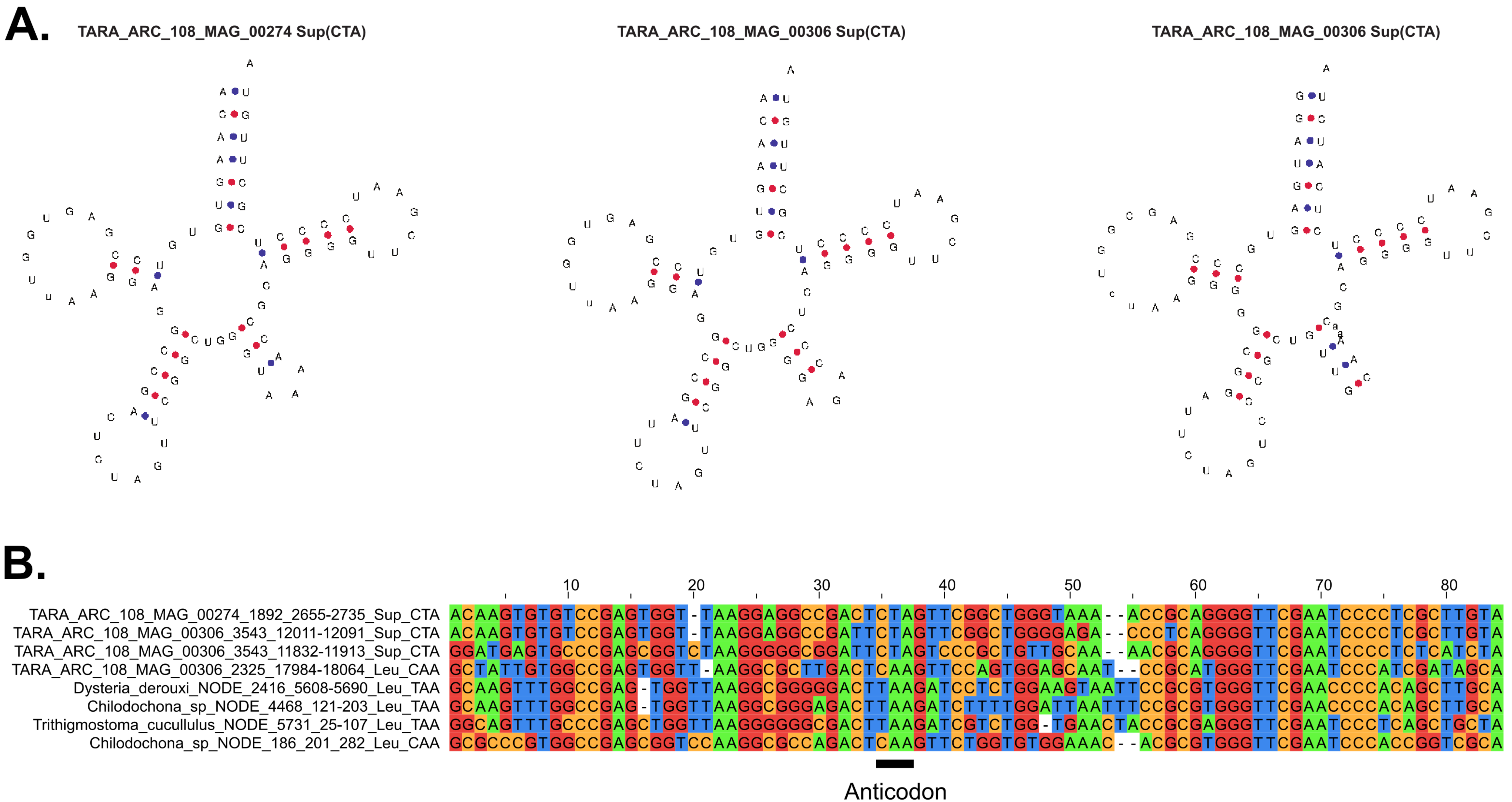
